# Deep characterization of the protein lysine acetylation in human gut microbiome and its alterations in patients with Crohn’s disease

**DOI:** 10.1101/772483

**Authors:** Xu Zhang, Zhibin Ning, Janice Mayne, Shelley A. Deeke, Krystal Walker, Charles L. Farnsworth, Matthew P. Stokes, David Mack, Alain Stintzi, Daniel Figeys

## Abstract

Metagenomic and metaproteomic approaches have been used to study the composition and functions of the microbiota. However, no studies have examined post-translational modifications (PTM) on human microbiome proteins at the metaproteome level, and it remains unknown whether the microbial PTM is altered or not in patient microbiome. Herein we used anti-acetyl-lysine (Kac) antibody enrichment strategy and mass spectrometry to characterize the protein lysine acetylation in human microbiome, which successfully identified 35,200 Kac peptides corresponding to 31,821 Kac sites from the microbial or host proteins in human gut microbiome samples. The gut microbial proteins exhibited Kac motifs that were distinct from those of human proteins. Functional analysis showed that microbial Kac proteins were significantly enriched in energy production and abundant in enzymes related to transferases and oxidoreductases. Applying to the analysis of pediatric Crohn’s disease (CD) patient microbiome identified 52 host and 136 microbial protein Kac sites that were differentially abundant in CD versus controls. Interestingly, most of the decreased Kac sites in CD were derived from Firmicutes and most of the increased sites were derived from Bacteroidetes. Forty-six out of the 52 differentially abundant human protein Kac sites were increased in CD patients, including those on calprotectin, lactotransferrin and immunoglobulins. Taken together, this study provides an efficient approach to study the lysine acetylation in microbiome and revealed taxon-specific alterations in the lysine acetylome as well as changes in host protein acetylation levels in intestinal samples during the on-set of disease in CD patients.

## Introduction

The intestinal microbiome is emerging as an important “organ” within the human body that actively interacts with its host to influence human health [1]. Dysbiosis of the intestinal microbiota has been reported to be associated with a myriad of diseases, including obesity, diabetes, Crohn’s disease (CD), cancer, and cardiovascular diseases [2]. In the past few years, meta-omic approaches, including metagenomics, metatranscriptomics and metaproteomics, have been applied to study the alterations of the microbiome composition and functions in patients with these diseases [3–6]. However, very little is known of the regulatory processes in the microbiome, such as post-translational modifications (PTMs) that are known to regulate the activity of proteins. In fact, there are currently no published studies on the global and deep characterization of PTMs in the human microbiome, nor published techniques for the efficient PTM profiling at the metaproteome level.

Acetylation is an important PTM in both Eukaryotes and Prokaryotes [7, 8]. In particular, lysine (N_ε_) acetylation has been shown to be involved in the regulation of various biological processes, including transcription and metabolism [8, 9]. Protein lysine acetylation has been characterized in several single bacterial species including *Escherichia coli* [10–13], *Bacillus subtilis* [14], *Salmonella enterica* [8] and *Mycobacterium tuberculosis* [15]. In bacteria, up to 40% of proteins can be acetylated [16]. Lysine acetylation has been widely implicated in various microbial processes including chemotaxis [17], nutrient metabolism [13], stress response [13] and virulence [18]. These findings highlight the importance of studying protein acetylation in the microbiome. However, as mentioned above, no study has yet globally examined protein acetylation in human microbiomes, and very little is known about protein acetylation within the microbiome in the context of diseases such as CD.

In this study, we first established experimental and bioinformatic workflows for characterization of the microbiome lysine acetylation. Briefly, an immuno-affinity-based approach was used for the enrichment of acetyl-lysine (Kac) peptides from the microbiome protein digests; the eluted peptides were analyzed with Orbitrap-based mass spectrometer (MS); and the MS data was then processed using an integrative metaproteomics/lysine acetylomics bioinformatic workflow that was developed in the current study. Lysine acetylome profiling of the human gut microbiome was then performed with fecal samples collected from six healthy adults. In total, 35,200 Kac peptides corresponding to 31,821 Kac sites were identified from either human or microbial proteins, which enabled deep characterization of the Kac motif and protein functions in the human microbiome. To the best of our knowledge, this is the first global characterization of Kac proteins in human microbiomes and achieved the highest number of site identification in lysine acetylomic studies. We further applied the approach to study the alterations of lysine acetylation in intestinal aspirate samples collected from the descending colon mucosal-luminal interface (MLI) of children with new-onset Crohn’s disease (CD) or controls. This analysis demonstrated the upregulation of Kac in host proteins, such as immune-related proteins, and down-regulation of Kac in microbial proteins from the Firmicutes species that are known short-chain fatty acid (SCFA)-producers, in CD compared with controls. This study provides an efficient workflow for studying lysine acetylation in the microbiomes and our results provide additional information on the intestinal dysbiosis in pediatric CD.

## Results

### Integrative metaproteomic/lysine acetylomic workflow enable deep identification of Kac peptides in microbiome

In this study, the proteolytic peptides generated from each microbiome sample were divided into two aliquots to allow for both metaproteomics and lysine acetylomics analysis. Lysine acetylated peptides from the first aliquot were enriched using a seven-plex anti-Kac peptide antibody cocktail (PTMScan® acetyl-lysine motif [Ac-K] kit) [19]; the second aliquot was directly analysed for metaproteome profiling (Fig. 1a). An integrative metaproteomics/lysine acetylomics data processing workflow was then developed based on our previously established MetaPro-IQ workflow [20] and MetaLab software tool [21] (Fig. 1b). Briefly, each of the raw files was first searched against the integrated gut microbial gene catalog (IGC) protein database using MetaLab [21]; the parameters were as default except that lysine acetylation (m/z 42.010565, H[2]C[2]O) was added as an additional variable modification. The sample-specific databases for both aliquots of all samples were then combined and redundant proteins removed. The resulting reduced database was then concatenated with a human protein database for peptide/protein identification and quantification of both metaproteome and lysine acetylome data sets.

**Figure 1.**
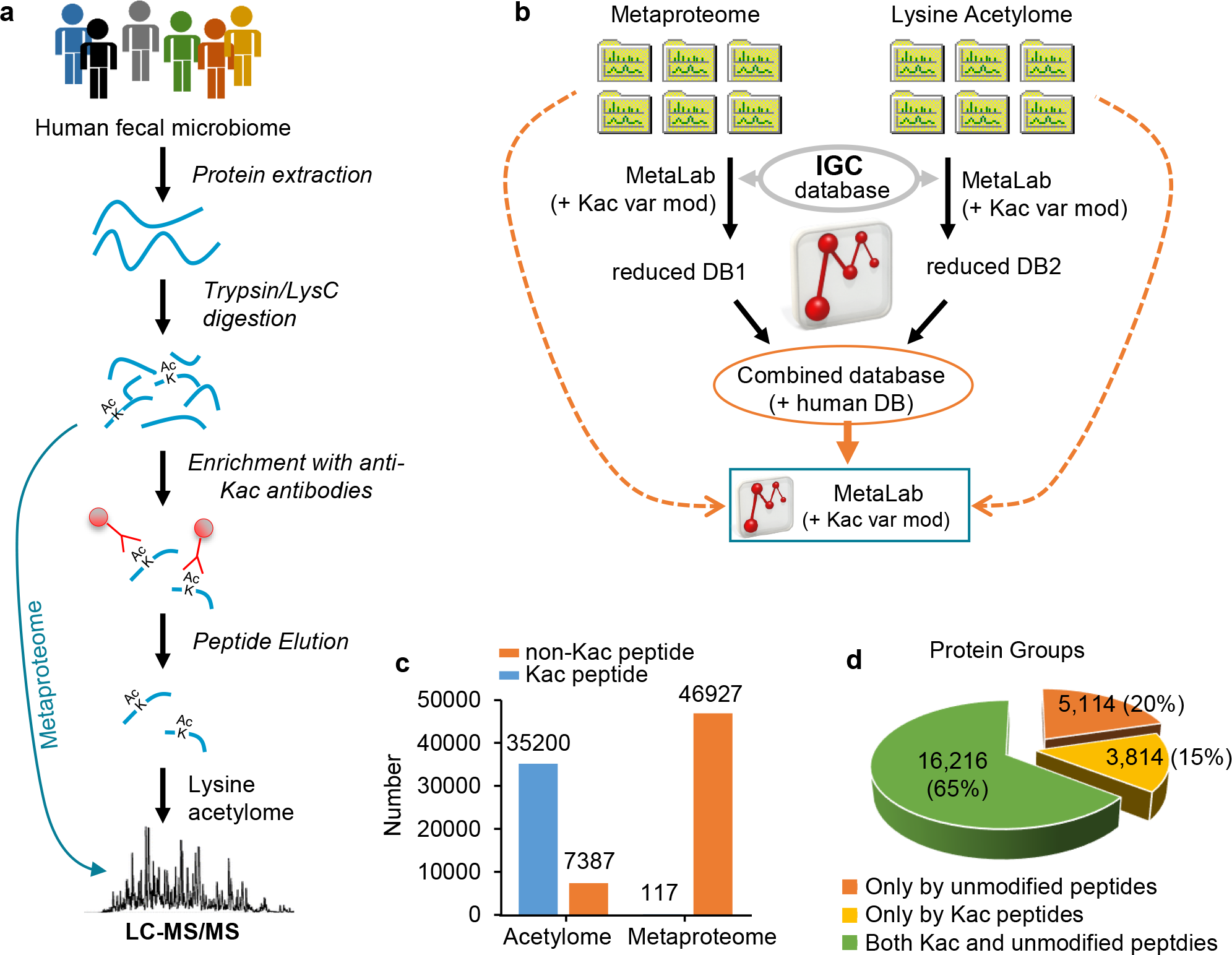
Experimental and bioinformatic workflows for integrative metaproteomic and lysine acetylomic characterization of the microbiome. (a) Experimental workflow; (b) Integrative metaproteomics/acetylomics data processing workflow; (c) Total number of identified Kac and non-Kac peptides in metaproteomic and lysine acetylomic aliquots, respectively; (d) Distribution of identified protein groups with the non-Kac peptide and Kac peptide sequences in the whole data set (both lysine acetylomic and metaproteomic aliquots).

This integrative bioinformatic workflow enabled the generation of comparable quantitative results at protein and Kac levels since the same database was used for both data sets. In total, this study identified 46,927 non-Kac peptides and 117 Kac peptides (171 Kac sites) from the metaproteomic aliquot; in contrast, 35,200 Kac peptides (31,821 Kac sites) and 7,387 non-Kac peptides were identified from the lysine acetylomic aliquot (Fig. 1c). This result indicates a high efficient enrichment of Kac peptides with the anti-Kac antibody cocktail, namely 83% of the identified peptides were Kac peptides. Among the 31,821 Kac sites identified, 1662 sites were quantified in all six samples and 6206 in at least 4 samples (Supplementary Fig. S1). Among the 117 Kac peptides identified in metaproteomic aliquot, only 6 were also identified in lysine acetylomic aliquot. To evaluate the overlap of identified Kac proteins with proteins identified in unenriched samples, we performed combined database search with raw files from both lysine acetylomic and metaproteomic aliquots. In total, this study identified 25,144 protein groups with 20,030 (80%) having at least one Kac modification. Interestingly, there were 3814 protein groups (15%) that were only inferred from Kac modified peptides (Fig. 1d), suggesting an efficient enrichment of low abundant Kac proteins/peptides using the current enrichment approach.

### Characterization of the host and microbial Kac motifs in the human gut microbiome samples

For the characterization of Kac sites, we used the 31,821 Kac sites identified in lysine acetylomic aliquot, of which 31,307 were from microbes and 497 were of human origin (Fig. 2a). We first characterized the amino acid distribution surrounding the acetylated lysine, for both human and microbial Kac sites (Fig. 2a). Both human and microbial Kac sites showed high frequency of glutamic acid (E) at −1 and leucine (L) and +1 positions. The microbiome protein Kac sites were frequently flanked by repeats of the small, hydrophobic amino acid, alanine (A); this was less common for the human protein Kac sites (Fig. 2a). We further analyzed and visualized Kac protein motifs using pLogo [22] in a sequence window of 13 amino acids (Fig. 2b and c). For human Kac sites, significant over-representation was only observed for tyrosine (Y) at position −4 and +1 (Fig. 2b and Supplementary Data 1). In contrast, 68 significantly over-represented events were observed for microbiome protein Kac sites, with E at position −1 and phenylalanine (F) at position +1 being the most significantly over-represented (Fig. 2c and Supplementary Data 1). Interestingly, small amino acid, alanine, was observed as significantly over-represented at all the 12 positions assessed and its occupancy frequency was >9% (10.3 ± 0.7%; mean ± SD) for all positions (Supplementary Data 1). In contrast, the median occupation frequency of all other significantly over-represented events was 5.8% (5.8 ± 3.6%; mean ± SD). These observations further confirm the high frequency of alanine near the Kac sites in microbiome proteins.

**Figure 2.**
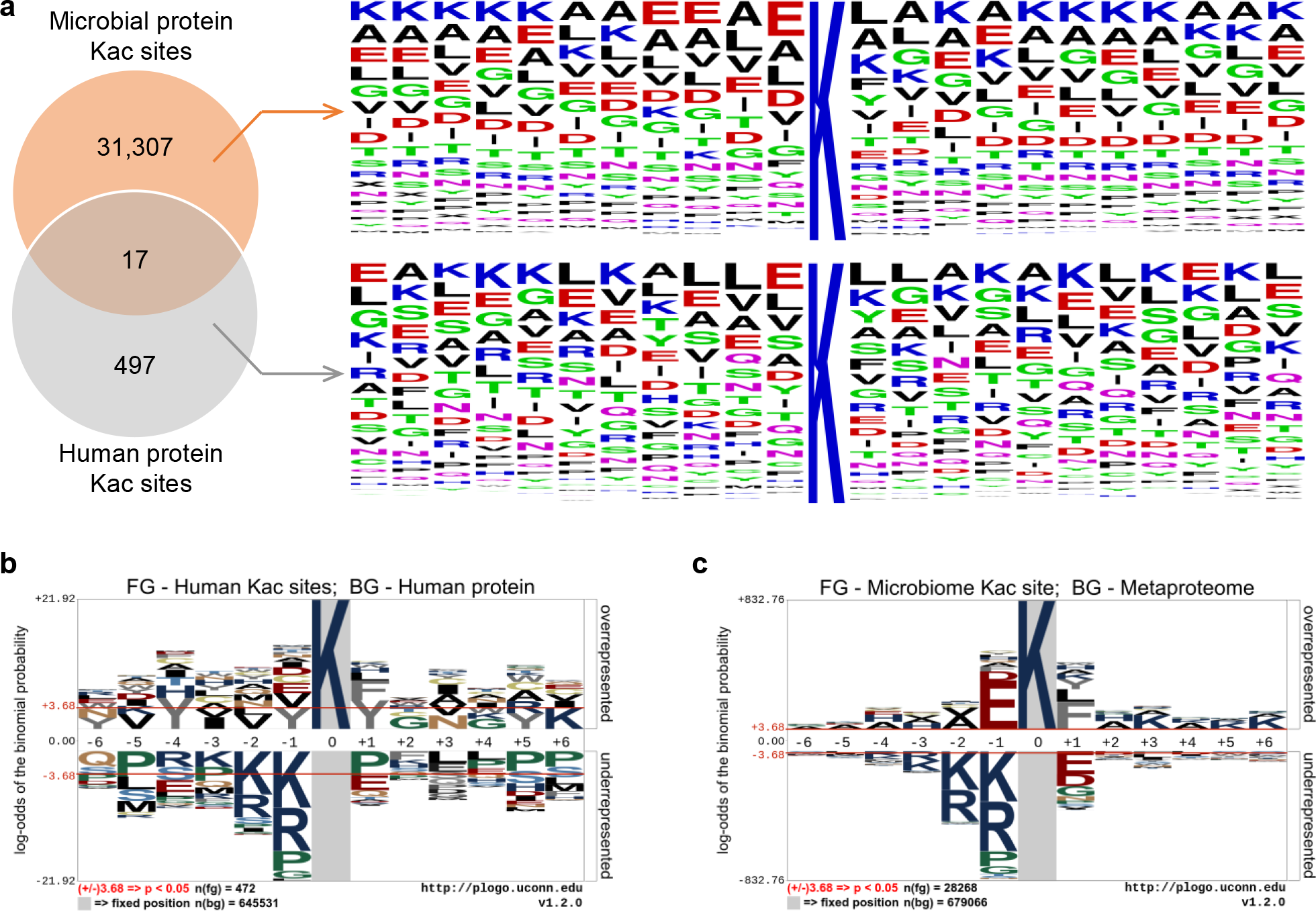
Characterization of the identified Kac sites in the microbial and human proteins from human fecal microbiome samples. (a) Amino acid composition analysis of identified human and microbial Kac peptide sequences. Venn diagram shows the overlap of identified human and microbial protein Kac sites; (b-c) pLogo sequence logo visualizations of all identified human (b) and microbiome (c) Kac sites. The n(fg) and n(bg) values indicate the number of foreground and background sequences, respectively. The red horizontal bars on the pLogo correspond to a threshold of p < 0.05.

We then examined whether different bacteria showed different protein Kac motifs in microbiome. Among the five bacterial phyla (Firmicutes, Bacteroidetes, Actibobacteria, Proteobacteria and Fusobacteria) with >100 Kac sites identified in this data set, high similarity between different phyla was observed (all showed high frequency of E at position −1 and F at position +1; Supplementary Fig. S2a-e). Firmicutes and Bacteroidetes are the two most abundant bacterial phyla in human gut microbiota and obtained the highest number of Kac sites identified. Motif analysis using pLogo identified 57 significantly over-represented events (position-amino acid pairs) for Firmicutes and 41 significantly over-represented events for Bacteroidetes. Interestingly, 35 out of the 41 significantly over-represented events in Bacteroidetes were also significantly over-represented events in Firmicutes (Supplementary Fig. S2f), indicating high conservation of Kac motif in different human gut microbial species.

### Taxonomic and functional characterization of microbial Kac proteins in the human gut microbiome

To characterize the taxonomic and functional distribution of microbial Kac proteins, we uploaded all the 35,200 Kac peptide sequences identified in this study into Unipept for biodiversity analysis [23]. Among all the Kac peptide sequences, 28,321 peptides (80%) were assigned to the kingdom Bacteria and 24,785 peptides could be classified at phylum level (15,170 from Firmicutes, 7876 from Bacteroidetes and 1739 from other phyla; Fig. 3a and Supplementary Data 2). A high proportion of the Kac peptides were from bacteria belonging to four genera: *Prevotella*, *Faecalibacterium*, *Bacteroides*, *and Eubacterium* (Supplementary Data 2). We calculated the Firmicutes-to-Bacteroidetes (F/B) ratios based on the intensities of their distinctive peptides yielding an average of 6.35 in lysine acetylome, which was significantly higher than that of metaproteome (an average of 4.90; paired T-test, P = 0.04).

**Figure 3.**
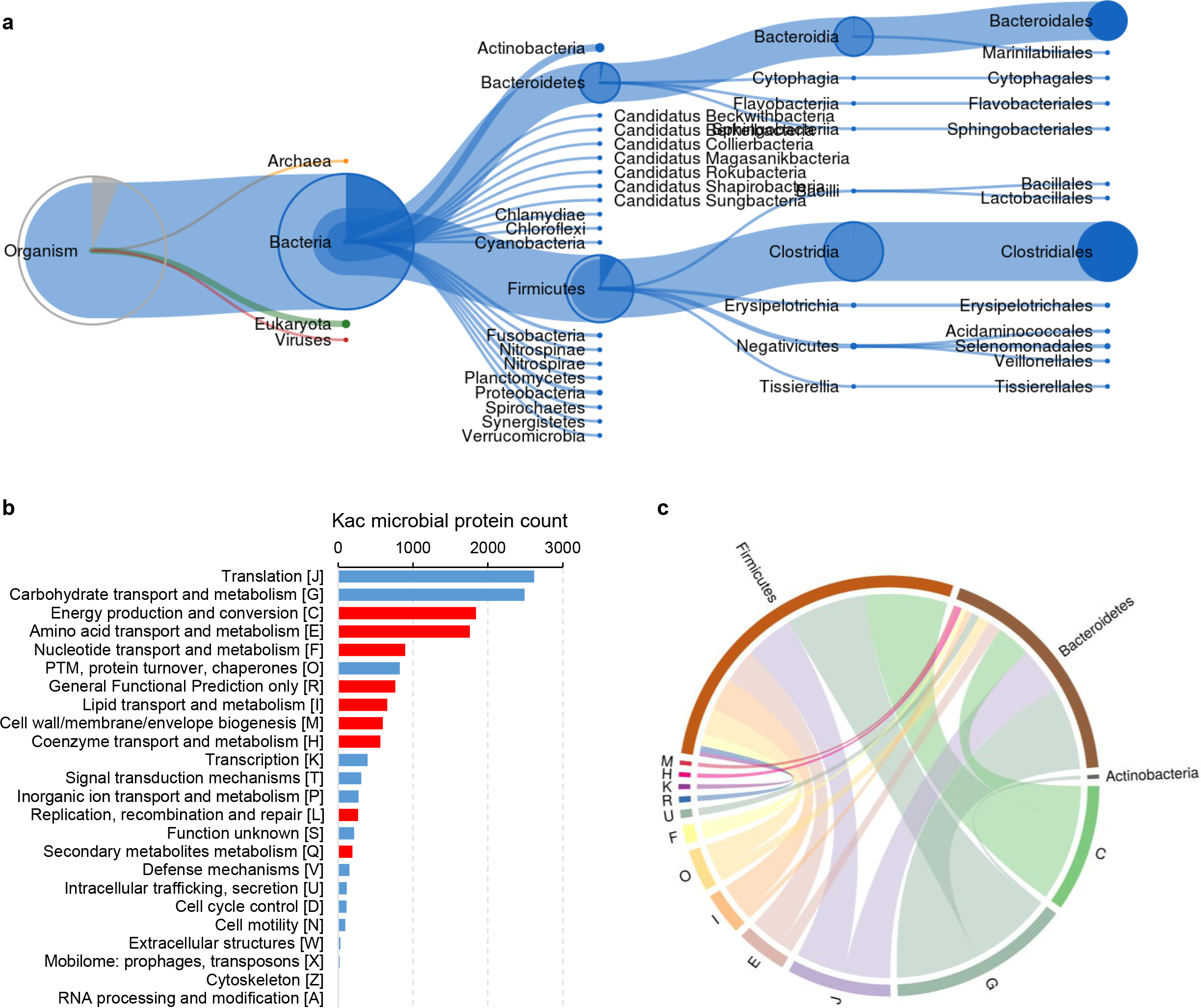
Taxonomy and functions of the identified Kac proteins in human fecal microbiome samples. (a) Treeview plot of microbial taxa that were assigned using all of the identified Kac peptides. Treeview plot were generated using Unipept (https://unipept.ugent.be/). (b) COG category distribution of microbial Kac proteins. Significantly enriched categories are highlighted in red. Significance was determined with a hypergeometric test using the unmodified microbial proteins identified in the metaproteomic samples as background. (c) Taxon (phylum)-specific functions of identified microbial Kac proteins. Circos plot were generated using iMetaLab tool suites (https://imetalab.ca/).

Gene ontology (GO) term annotation of the Kac peptides showed that the top GO biological processes were translation (2642 peptides) and carbohydrate metabolism (1660 peptides); the top GO molecular functions were ATP binding (6190 peptides) and metal ion binding (3439 peptides); and top GO cellular components were cytoplasm (7190 peptides) and ribosome (2102 peptides) (Supplementary Data 3). Enzyme Commission (EC) number annotation yielded a total of 1025 enyzmes from 5 enzyme classes (Supplementary Data 4). The most Kac modified enzymes identified in this study belonged to class EC 2 (transferases; mainly EC 2.7: transferring phosphorus-containing groups) and EC 1 (oxidoreductases).

We then performed functional enrichment analysis for microbial Kac proteins with the Clusters of Orthologous Groups (COG) database. Using unmodified microbial proteins identified in metaproteomic aliquot as background, the enrichment analysis revealed that microbial Kac proteins were significantly enriched in energy production and conversion, transport or metabolism of several molecules (including amino acid, nucleotide, lipid and coenzyme), cell wall biogenesis, replication as well as secondary metabolites metabolism (Fig. 3b). Taxon (phylum)-specific functional analysis further showed that Kac proteins from Firmicutes contributed to the majority proteins in energy production and conversion, and those from Bacteroidetes contributed to nearly half of the Kac proteins in translation and carbohydrate transport/metabolism although their total abundances were much lower than those from Firmicutes (Fig. 3c).

### Alterations of the lysine acetylome in the intestinal microbiome of pediatric CD patients

To demonstrate the applicability of the integrated lysine acetylomic/metaproteomic approach, we then analysed 18 intestinal aspirate samples collected from 10 pediatric CD patients and 8 control subjects. The average age of the CD patients was 13 ± 2 years (mean ± SD; 6 male/4 female) and the average age of control patients was 13 ± 5 years (mean ± SD; 2 male/6 female). The CD patients included 4 with mild disease (mean PCDAI = 18; SD = 4), 6 with moderate/severe disease (mean PCDAI = 41; SD = 7) (Supplementary Data 5). In total, we accurately quantified 4623 Kac sites in lysine acetylomic aliquot and 17,684 protein groups in metaproteomic aliquot. Principal component analysis (PCA) showed a trend to separate both the metaproteome and the lysine acetylome for CD versus control (Fig. 4a and b). Partial least square discriminant analysis (PLS-DA) of the metaproteome (Q^2^ = 0.59) identified 208 increased and 284 decreased protein groups in CD compared to control subjects. PLS-DA of the lysine acetylome (Q^2^ = 0.33) identified 51 increased and 29 decreased Kac sites in CD compared to control subjects. In addition to those PLS-DA-identified differentially abundant proteins or Kac sites, the proteins and Kac sites that were detected in greater than 80% of samples in the CD group and less than 20% of samples in the control group were also considered as increased in CD compared to controls. Similarly, the proteins and Kac sites that were detected in less than 20% of samples in CD group and greater than 80% of samples in control group were considered as decreased in CD. Altogether, the current study identified 82 Kac sites that were increased, and 106 Kac sites that were decreased, in CD compared to control subjects (Supplementary Data 6).

**Figure 4.**
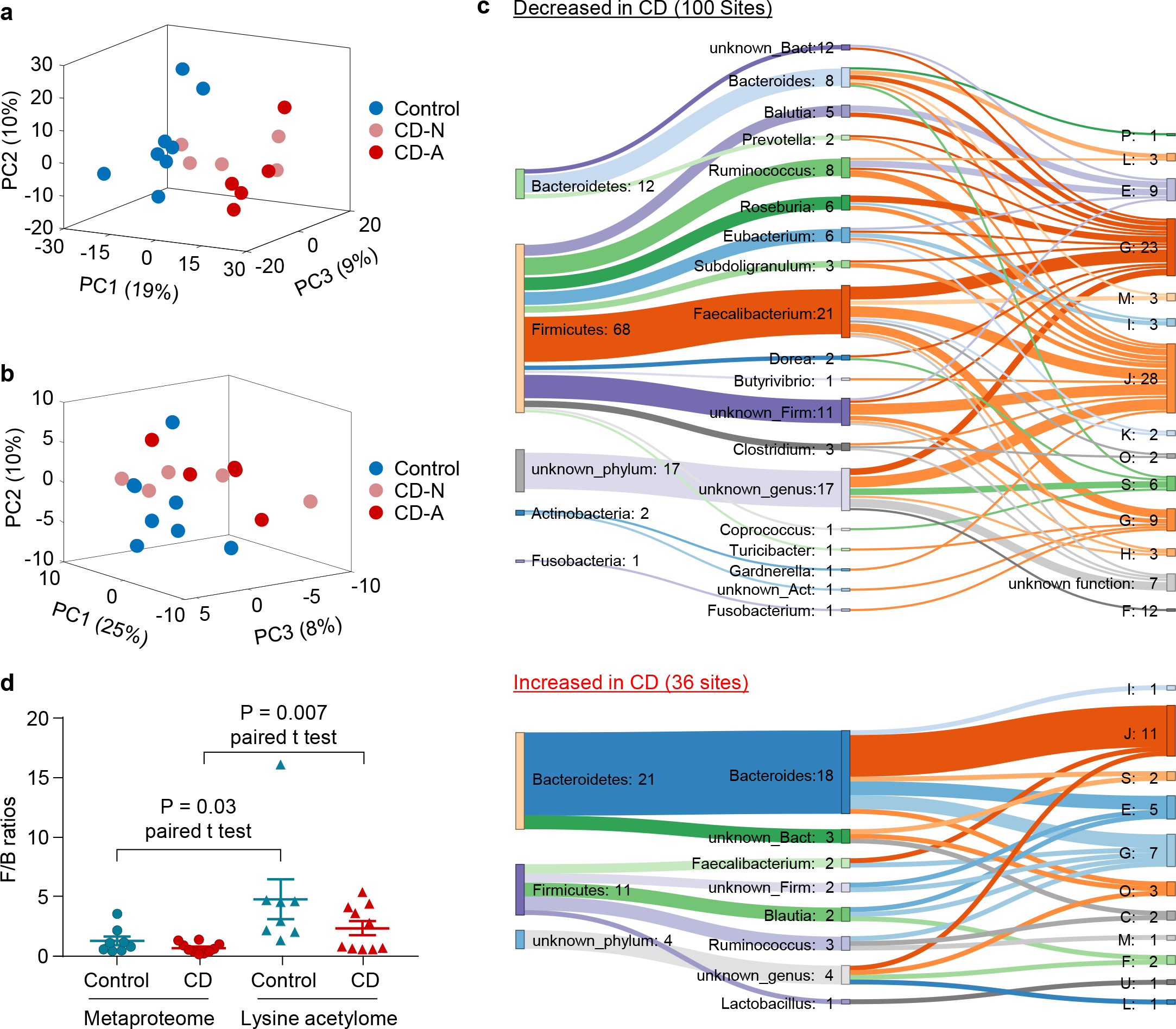
Lysine acetylome alterations of the intestinal microbiome in pediatric CD. (a) PCA score plot of the metaproteome of the intestinal microbiome. (b) PCA score plot of the lysine acetylome of the intestinal microbiome. (c) Differentially abundant microbial Kac sites. The COG category and taxonomy (phylum and genus) for the differentially abundant Kac sites are shown in the Sankey plot. The numbers after the colon “:” indicate the numbers of differentially abundant Kac sites. The phylum-genus links and genus-function (COG category) links are shown. Each letter corresponds to a COG category as shown in Figure 3. (d) Firmicutes-to-Bacteroidetes (F/B) ratios calculated using the lysine acetylome- or metaproteome-based abundances.

Among the 82 Kac sites that were increased in CD compared to control subjects, 46 were from human proteins and 36 were from microbiome proteins. However, only 6 Kac sites that were decreased in CD were from human proteins and the remaining 100 upregulated Kac sites were from microbiome proteins. Interestingly, 68 out of the 100 upregulated microbial Kac sites were derived from Firmicutes, and 21 out of the 36 down-regulated microbial Kac sites were derived from Bacteroidetes (mainly from the genus *Bacteroides*) (Fig. 4c). The majority of the 68 upregulated Firmicutes-derived Kac sites were from known SCFA-producing bacteria, including *Faecalibacterium* (21 sites), *Ruminococcus* (8 sites), *Eubacterium* (6 sites), *Roseburia* (6 sites) and *Blautia* (5 sites). Taxon-specific functional analysis showed that the microbial Kac sites that showed increased abundances in CD were mainly from translation-related proteins of *Bacteroides*; and the down-regulated microbial Kac sites in CD were mainly from proteins that are involved in translation and carbohydrate metabolism of known SCFA-producers as mentioned above (Fig. 4c). For example, among the 21 protein Kac sites of *Faecalibacterium* that were decreased in CD, 6 sites were from proteins related to carbohydrate metabolism, 4 from proteins related to energy metabolism, and 5 from proteins related to translation.

We also performed comparative taxonomic analysis using the quantified Kac microbial peptides and linear discriminant analysis effect size (LEfSe) analysis. The results showed that the acetylome-based abundances of species *Roseburia inulinivorans*, *Eubacterium eligens* and *Megamonas funiformis* were significantly decreased in CD compared to that of controls, and the abundance of *Bacilli* was significantly increased although its metaproteome-based abundance was decreased in CD (Supplementary Fig. S3 and Supplementary Note 1). The calculation of F/B ratios in both lysine acetylome and metaproteome showed that lysine acetylome displayed higher F/B ratios than metaproteome in both control (P = 0.03) and CD groups (P = 0.007) (Fig. 4d), which was in agreement with the above observations in adult fecal microbiomes in this study.

### Altered abundances of Kac sites on human proteins that are associated with intestinal microbiomes in pediatric CD

As mentioned above, we identified 46 human protein Kac sites that showed increased abundances and six human protein Kac sites that showed decreased abundances in CD compared to control (Fig. 5). The six decreased human protein Kac sites in CD were on chymotrypsin-like elastase family member 3A (CEL3A; 2 sites), CEL3B (2 sites), ubiquitous mitochondrial creatine kinase (KCRU) and Xaa-Pro aminopeptidase 2 (XPP2). The increased human protein Kac sites were mainly from the proteins lactotransferrin (LTF; 10 sites) and calprotectin (protein S100A8 and S100A9; 7 sites). In addition, there were 11 elevated protein Kac sites from immunoglobulins (Ig), including Ig heavy constant alpha (IGHA1 and IGHA2), Ig heavy constant mu (IGHM), Ig lambda constant 3 (IGLC3), Ig kappa constant (IGKC) and Ig lambda-like polypeptide 5 (IGLL5). These findings indicate a potential role of protein acetylation in the aberrant immune responses in the intestine of pediatric CD patients.

**Figure 5.**
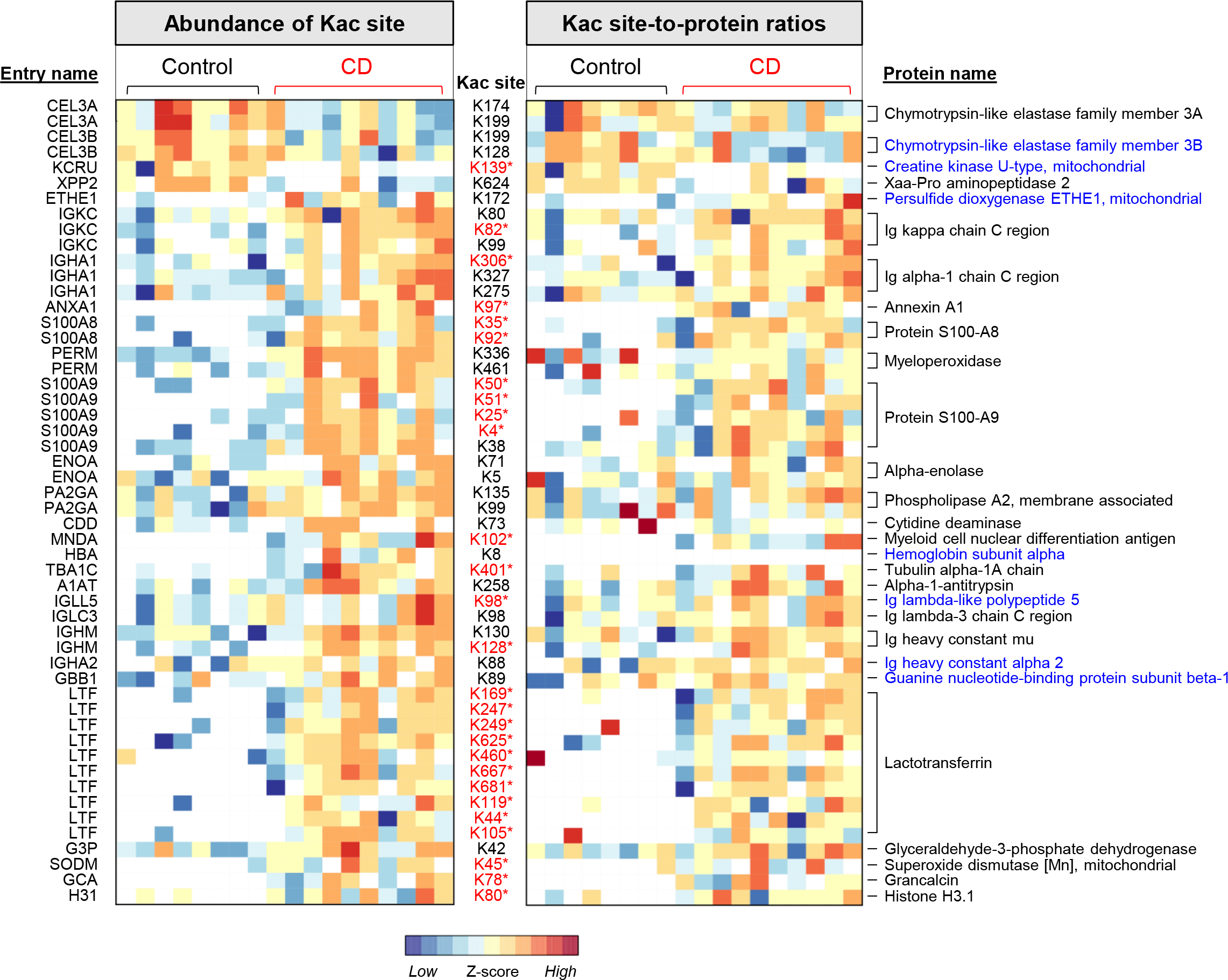
Abundance alterations of human protein Kac sites in the intestinal microbiome samples in CD. A heatmap of differentially abundant human protein Kac sites is shown on the left and a Kac site-to-protein ratio heatmap is shown on the right. Each row of the heatmap is a protein Kac site (indicated in between the two panels). The UniProt protein entry name and protein name for each Kac site are indicated on the left side and right side, respectively. The Kac sites highlighted in red (*) retained the differences in their site-to-protein ratios. Protein names highlighted in blue indicate the proteins with no significant difference between CD and control in unenriched samples.

To study the relative abundances of Kac sites to their total proteins, we calculated the ratios between each of the 52 differentially abundant Kac sites and their corresponding protein abundances in the metaproteomic aliquot (Fig. 5). Twenty-seven out of the 52 sites exhibited significantly different site-to-protein ratios (Mann–Whitney test, P < 0.05) or frequencies of detection between CD and control subjects. In total, the 52 differentially abundant human protein Kac sites mapped to 26 proteins. Of these, six proteins (CEL3B, KCRU, persulfide dioxygenase, IGLL5, IGA2 and G protein subunit beta 1) were not significantly different at the protein abundance level between CD and control subjects, and one protein (hemoglobin subunit alpha; HBA1) was not detected in the metaproteomic analysis.

In the current study, we found that the total protein levels of both S100A8 and S100A9, two monomers of a known CD biomarker calprotectin [24, 25], were significantly increased in CD compared to controls (Supplementary Fig. S4). In addition, we identified 5 Kac sites for protein S100A8 and 7 Kac sites for protein S100A9 in the lysine acetylome data set, among which 2 for S100A8 and 5 for S100A9 were detected in <20% of control samples and >80% of CD samples (Fig. 6a-b). Among all the identified calprotectin Kac sites, only one site for S100A8 (K18) and one site for S100A9 (K38) were quantified in ≥3 control samples, and both showed significantly lower abundances in control than CD (Fig. 6a-b). Although there is no significant difference on the site-to-protein ratios of individual Kac site between CD and control samples (Supplementary Fig. S5), the ratios between the sum intensities of all Kac sites on S100A8, S100A9 and their corresponding proteins were significantly higher in CD compared to control samples (S100A8, P = 0.02; S100A9, P = 0.02; S100A8/9, P = 0.01) (Fig. 6c).

**Figure 6.**
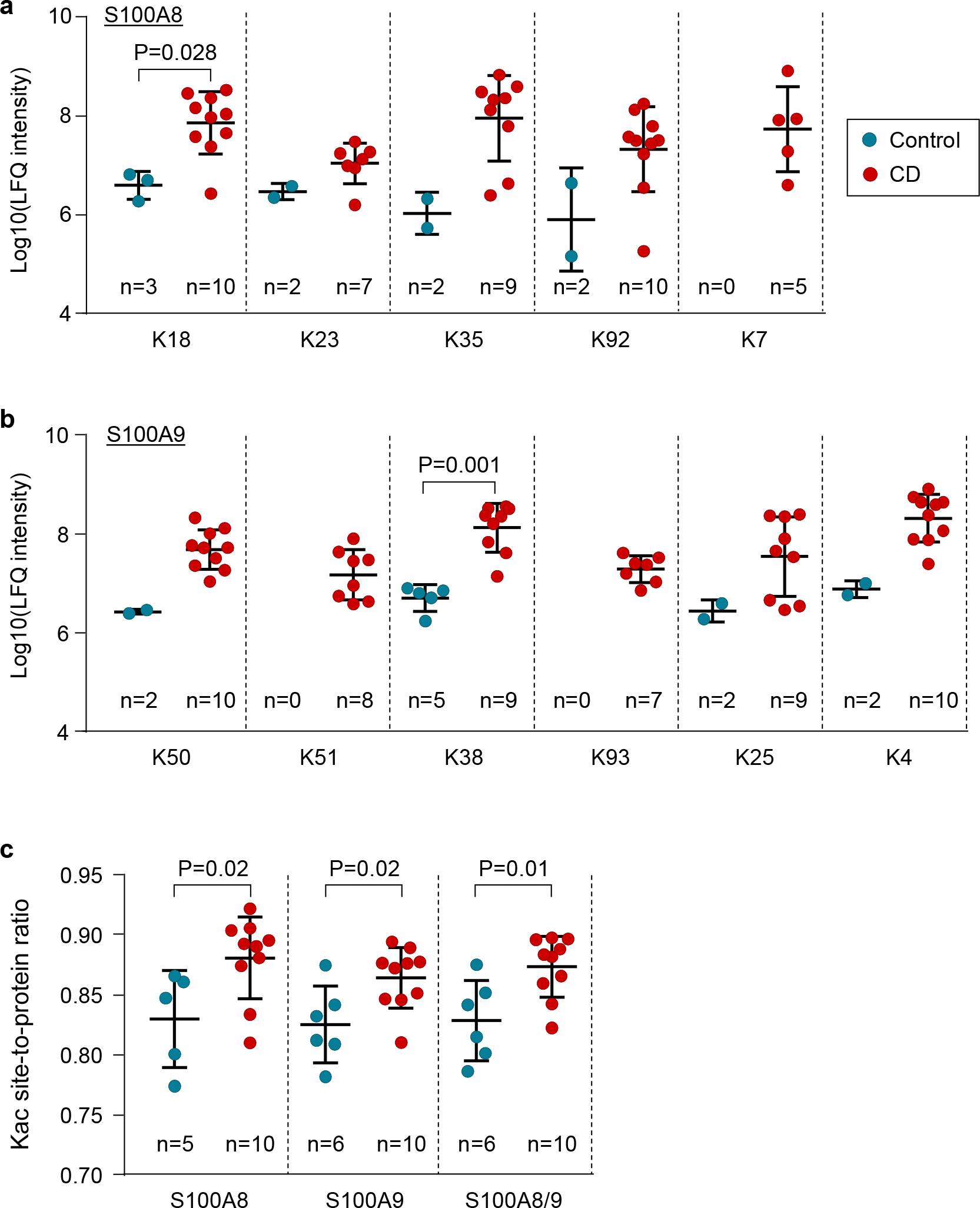
Changes in lysine acetylation of calprotectin in the intestinal aspirate samples of pediatric CD patients. (a) Kac sites of protein S100A8; (b) Kac sites of protein S100A9. K54 were not shown because they were quantified in only 1 sample. (c) Relative abundance of total lysine acetylation of S100A8 or S100A9. Ratios between the sum Kac site intensity (log10-transformed) and that of their corresponding protein in unenriched aliquot were shown.

## Discussion

In the current study, we successfully utilized a recently developed anti-Kac motif antibody mixture [19] for the enrichment of both human and microbial Kac peptides from human gut microbiome samples. We demonstrated that the microbial Kac proteins were significantly enriched in pathways such as energy production and lipid metabolism, which is in line with the consensus that protein lysine acetylation plays ubiquitous roles in both translation and metabolism [8, 9]. We also identified Kac motifs in microbiome proteins that were distinct from that of human proteins. We examined both the host and microbiome lysine acetylomes in intestinal samples collected from pediatric CD patients and controls, and found elevated Kac abundances of host immune response proteins in CD. Moreover, the abundances of Kac sites of proteins from known SCFA-producers were found to be decreased in CD compared with controls. These findings indicate a potential role of lysine acetylation on both host and microbial proteins in the dysbiotic host-microbiome interactions during the onset of CD.

Lysine acetylation is an important PTM event regulating various biological processes and cellular functions in all kingdoms of organisms [7, 8, 26, 27]. The global profiling of Kac has been performed in many organisms, including bacterial species such as *Escherichia coli* [10–13]. However, as mentioned above, the global characterization of Kac sites in the microbiome has not yet been studied. This is mainly due to the extremely high complexity of microbiome samples, which requires an enrichment approach with better coverage, and due to the bioinformatic challenges in efficiently identifying and quantifying the microbiome Kac peptides. In the current study, we utilized the seven anti-Kac monoclonal antibody cocktail, which was developed by Svinkina *et al.* [19], for the enrichment of Kac peptides from tryptic digest of microbiome proteins. In addition, we developed an integrated metaproteomics/lysine acetylomics data processing workflow based on our previously developed MetaPro-IQ approach [20] and MetaLab software tool [21]. Altogether, the experimental and bioinformatic workflow enabled a successful Kac peptide enrichment, identification and quantification for microbiome samples. In total, over 35,000 Kac peptides were identified, which was 80% of the all identified peptides in enriched samples. This is far higher than that in unenriched samples (0.2% with only 117 Kac peptides). It is worth noting that only six out of the 117 Kac peptides that were identified in unenriched samples overlapped with those identified in the enriched samples. This finding indicates that the Kac peptides identified from the unenriched aliquot were likely false identifications and suggests that an enrichment step may be necessary to reliably identify PTMs in the microbiome.

Firmicutes is one of the most abundant bacterial phyla in the human gut microbiota and plays important roles in human health at least in part through generating SCFAs and harvesting energy from indigestible dietary fibres [28, 29]. In the analyses of MLI microbiome from pediatric CD patients, we identified 1710 Kac peptides corresponding to 1258 protein groups from Firmicutes, which was nearly twice the number of Kac peptides from Bacteroidetes (999 Kac peptides corresponding to 836 protein groups). Accordingly, the average F/B ratios were significantly higher in the lysine acetylome than the metaproteome for both control (4.8 vs. 1.3) and CD microbiomes (2.4 vs. 0.7). These findings suggest that Firmicutes may have a higher protein Kac levels than Bacteroidetes in the human gut. Many species in Firmicutes are known producers of butyrate and acetate, such as species from *Faecalibacterium*, *Blautia* and *Roseburia* [30–33]. Butyrate is a known inhibitor for lysine deacetylase (KDAC) [34] and acetate can be used to generate acetyl-CoA, a substrate and acetyl donor for lysine acetyltransferase (KAT)-mediated lysine acetylation [13]. In fact, many proteins involved in acetate metabolism, including acetyl-CoA synthetase which converts acetate to acetyl-CoA, are acetylated proteins and their activities are also regulated by lysine acetylation [13]. Therefore, the potential regulating effects of these microbial metabolites (i.e., butyrate and acetate) may lead to higher global protein acetylation levels in SCFA-producers than others. Although further mechanistic studies are still required, our findings suggest that lysine acetylation might be a potential target for manipulating the growth and functional activity of SCFA-producers for the treatment of diseases.

Intestinal dysbiosis, in particular the depletion of SCFA-producing bacteria, is commonly associated with the development of both adult and pediatric CD [35–37]. In this study, we demonstrated that the intestinal dysbiosis observed in pediatric CD patients includes alterations in the lysine acetylation of microbiome proteins, in particular the proteins of butyrate/acetate-producers such as *R. inulinivorans*, *E. eligens* and *M. funiformis* (Figure 5F). Further studies are needed to understand whether the decrease of protein Kac levels in butyrate/acetate-producers in CD patients may contribute to the depletion of SCFA-producers or not. Previous studies have shown that various KDAC inhibitors, such as butyrate, suberyolanilide hydroxamic acid (SAHA), valproic acid (VPA) and statin hydroxamate, effectively treat colitis in animal models [38–41]. In particular, Wei *et al.* reported that statin hydroxamate alleviated colitis and reduced the blood endotoxin (lipopolysaccharide) level [39], an indicator of dysbiosis of gut microbiota [42]. Although the mechanism has been attributed to their beneficial effects on the intestinal epithelium cells, our findings indicate that the KDAC inhibitors may in part directly interact with the microbiome for the alleviation of intestinal colitis.

## Conclusions

We demonstrated that a recently developed, commercially available anti-Kac monoclonal antibody cocktail can be used to successfully enrich Kac peptides from the human microbiome. Based on this, we established an experimental and bioinformatic data analysis workflow for microbiome-wide characterization of protein lysine acetylation. Analyzing the microbiome samples collected from the intestinal mucosal surface of pediatric CD and control subjects revealed that the alterations of the lysine acetylome of microbiome in pediatric CD patients were phylum-specific, with the majority of down-regulated Kac sites belonging to the SCFA-producing bacteria in Firmicutes. In addition to the microbiome Kac sites, we also identified Kac sites on human proteins that were associated with the microbiomes and demonstrated the alterations of Kac levels on immune response proteins, such as immunoglobulins and calprotectin. Our study provides a useful tool for the functional study of the microbiome at the PTM level and the findings provide additional information on the dysbiosis and onset of pediatric CD.

## Methods

### Subjects and sample collections

Fresh fecal samples were collected from six healthy adult volunteers at the University of Ottawa with protocol (Protocol # 20160585-01H) approval by the Ottawa Health Science Network Research Ethics Board at the Ottawa Hospital. MLI aspirate samples were collected from pediatric patients that were undergoing diagnostic colonoscopy for their medical problems and were suspected to possibly be CD. All participants (<18 years old) were treatment-naïve. CD was diagnosed through clinical, endoscopic, histologic and radiological evaluations according to standard criteria [43]. Disease severity was determined using the Pediatric Crohn’s Disease Activity Index (PCDAI) [44]. The Simplified-Endoscopy Score-Crohn’s disease (SES-CD) was used as segmental description of endoscopic mucosal characteristics (i.e. inflamed mucosa, amount of affected mucosa, ulcer size) [45]. Control patients had visually normal mucosa, histologically normal mucosal biopsies and normal imaging. The following exclusion criteria were implemented to further refine the cohort enrolled in this study: (1) presence of diabetes mellitus; (2) presence of infectious gastroenteritis within the past two months; (3) use of any antibiotics or probiotics within the past four weeks, or (4) irritable bowel syndrome. Subject clinical data were collected and managed using REDCap (Research Electronic Data Capture) hosted at the CHEO Research Institute. REDCap is a secure, web-based application designed to support data capture for research studies [46]. The MLI aspirate samples were collected as previously described [3, 35]. Briefly, any existing fluid and debris were first aspirated and discarded during colonoscopy. Sterile water was then flushed onto the mucosal surface to dislodge the mucus layer from the epithelial cells; the resulting fluid was then aspirated into a container. The latter was immediately put on ice and transferred to the laboratory for further processing.

### Sample processing, protein extraction and tryptic digestion

The fresh stool sample was immediately put on ice and subjected to differential centrifugation and wash according to the procedures described previously [47]. The resulting microbial pellets were then subjected to protein extraction using lysis buffer containing 4% (w/v) SDS, 50 mM Tris-HCl (pH 8.0) and protease inhibitor (cOmplete™, mini protease inhibitor cocktail; Roche Diagnostics GmbH). Protein lysates were then precipitated by adding five times the volume of lysis buffer of ice-cold acidified acetone/ethanol buffer overnight at −20 °C. Precipitated proteins were then collected with centrifugation at 16,000 g for 25 min at 4 °C, and washed three times by ice-cold acetone before re-suspending in 6 M urea, 100 mM ammonium bicarbonate buffer [3]. Protein concentration was determined using a DC protein assay (Bio-Rad Laboratories, Inc) according to the manufacturer’s instruction. Approximately 10~15 mg of proteins for each sample was then used for in-solution proteolytic digestion. Briefly, proteins were first reduced with 10 mM dithiothreitol (DTT) for 1 hour and 20 mM iodoacetamide (IAA) for 40 min at room temperature; then the samples were diluted by ten folds with 100 mM ammonium bicarbonate buffer followed by digestion using lysyl endopeptidase (Lys-C; Wako Pure Chemical Corp., Osaka, Japan) for 4 hours and trypsin (Worthington Biochemical Corp., Lakewood, NJ, USA) for overnight at room temperature. The resulting digests were then subjected to desalting using C18 column as described previously [47]. A small portion of the proteolytic peptides from each sample was used for metaproteomic analysis (directly load for MS analysis), and the remainder was used for Kac peptide enrichment.

For MLI aspirate samples, upon arriving at the laboratory, the samples were immediately mixed with protease inhibitor (cOmplete™, mini protease inhibitor cocktail; Roche Diagnostics GmbH). The aspirate samples were first centrifuged at 700 g for 5 min at 4 °C and the supernatant collected for another centrifugation at 14,000 g for 20 min at 4 °C. The pellet fraction was harvested for protein extraction using lysis buffer consisting of 4% (w/v) SDS, 8M urea, 50 mM Tris-HCl (pH 8.0) and cOmplete™ mini protease inhibitor cocktail. Protein lysates were then precipitated and washed using ice-cold acetone as described above. An equal amount (2.5 mg) of proteins for each sample was then used for in-solution trypsin digestion and desalting as described previously [47]. A small portion of the tryptic peptides (equivalent to 40 μg proteins) from each sample was used for metaproteomic analysis (directly load for MS analysis), and the remainder was used for Kac peptide enrichment.

### Kac peptide enrichment

Kac peptides were enriched using PTMScan^®^ Motif antibody kits (Cell Signaling technology, Inc.) according to the manufacturer’s instruction. Briefly, tryptic peptides were first re-suspended in PTMScan® IAP Buffer and centrifuged at 10,000 g for 5 min at 4 °C to remove any insoluble pellets. The supernatant was then added directly to the tube containing Kac motif antibody beads and mixed immediately by slowly pipetting up and down. The mixture was then incubated at 4 °C for 2 hr on a rotator. After incubation, the beads were collected by centrifugation at 2,000 g for 30 s. The beads were then washed twice with cold IAP buffer and three times with H_2_O. The peptides were eluted by adding 55 μl of 0.15% (v/v) trifluoroacetic acid (TFA) to the beads and incubating for 10 min while mixing gently. The supernatant was collected through centrifuging at 2,000 g for 30 s and the remaining beads were mixed with another 50 μl of 0.15% (v/v) TFA for another round of elution. Both eluents were then combined for desalting using C18 columns as described previously [47].

### Tandem mass spectrometry analysis

The peptides generated from metaproteomic and lysine acetylomic aliquotes of fecal microbiome samples were analyzed using Q Exactive HF-X mass spectrometer (ThermoFisher Scientific Inc.). For unenriched samples, peptides equivalent to 250 ng proteins were loaded for MS analysis; for enriched samples, all peptides were re-suspended in 20 μ 0.1% (v/v) formic acid, and 4 μ was used for MS analysis. Peptides were separated on an analytical column (75 μm × 15 cm) packed with reverse phase beads (1.9 μm; 120-Å pore size; Dr. Maisch GmbH, Ammerbuch, Germany) with 2 hr gradient from 5 to 35% (v/v) acetonitrile at a flow rate of 300 nl/min. The instrument method consisted of one full MS scan from 350 to 1400 m/z followed by data-dependent MS/MS scan of the 16 most intense ions and a dynamic exclusion duration of 20 s. Mass spectrometry analysis of MLI aspirate samples, including both enriched and unenriched samples, was performed on a Q Exactive mass spectrometer (ThermoFisher Scientific Inc.). For unenriched samples, peptides equivalent to 1 μg proteins were loaded for MS analysis; for enriched samples, all peptides were re-suspended in 20 μl was used for MS analysis. Briefly, peptides were μ separated on an analytical column (75 μm × cm) packed with reverse phase beads (1.9 μm; 120-Å pore size; Dr. Maisch GmbH, Ammerbuch, Germany) with 4 hr gradient from 5 to 35% (v/v) acetonitrile at a flow rate of 300 nl/min. The instrument method consisted of one full MS scan from 300 to 1800 m/z followed by data-dependent MS/MS scan of the 12 most intense ions, a dynamic exclusion repeat count of 2, and repeat exclusion duration of 30 s. The MS data were then exported in RAW format for further bioinformatic data processing.

### Identification and quantification of Kac and non-Kac peptides and proteins

Protein identification and quantification for both metaproteomic and lysine acetylomic data sets were performed using MetaLab [21] with a modified MetaPro-IQ workflow [20] as detailed in Figure 1B. In this study, we used the reference protein database that was generated in our previous metaproteomic studies of pediatric MLI aspirate samples [3], which we termed the IGC+ database in the current study. Briefly, the IGC+ database consists of protein sequences from human fecal microbial Integrated Gene Catalog (IGC) database [48], NCBI viral proteins, predicted protein sequences from shotgun metagenomic sequencing of MLI aspirate samples, and representative fungal species (details in [3]). The same database search parameters were used for both metaproteomic and lysine acetylomic data sets as follows: (1) up to four missed cleavages are allowed, (2) fixed modification includes cysteine carbamidomethylation, (3) potential modifications include methionine oxidation, lysine acetylation and protein N-terminal acetylation, and (4) a parent ion tolerance of 10 ppm and a fragment ion tolerance of 20 ppm. The peptide and protein identification were performed with a false discovery rate (FDR) threshold of 0.01.

The identified peptides and protein groups in unenriched samples were obtained from the “modificationSpecificPeptides.txt” and “proteinGroups.txt” files, respectively. The identified Kac peptides and sites in enriched samples were obtained from the “modificationSpecificPeptides.txt” and “Acetyl (K) Sites” files, respectively. Only Kac sites with a localization probability of >0.75 were used for further analysis.

### Kac motif analysis

The sequences of amino acids surrounding Kac sites were analyzed and visualized using WebLogo (https://weblogo.berkeley.edu/) [49]. Sequence windows of 11 amino acids surrounding Kac site were created. pLogo (https://plogo.uconn.edu/) [22] was used to identify statistically over-represented Kac motifs for the identified Kac sites. Briefly, sequence windows with six upstream and downstream amino acids surrounding the Kac site were extracted from the database and submitted to Motif-X for Kac motif extraction. The total identified microbial proteins from one randomly selected unenriched sample were used as background for microbial protein Kac motif extraction.

### Functional enrichment analysis

The COG category enrichment analysis of identified microbial Kac proteins was performed with hypergeometric probability analysis using the microbial proteins identified in unenriched samples as background. Briefly, each of the proteins identified in unenriched or enriched samples was annotated with a COG category as described previously [20]. The numbers of proteins assigned to each COG category from Kac proteins and background were used for calculating the significance P values of enrichment using *hygecdf* function and Benjamini-Hochberg adjusted FDR values using *mafdr* function in MATLAB (The MathWorks Inc.).

### Taxonomy and functional analysis

Taxonomic annotation of both unmodified and Kac peptides was performed using Unipept 4.0 with the “Equal I and L” and “Advanced misscleavage handling” options allowed [23]. Gene ontology (GO) term and Enzyme Commission (EC) number annotations were directly exported from Unipept analysis (https://unipept.ugent.be/). For enriched samples, only Kac peptides were used. Linear discriminant analysis (LDA) effect size (LEfSe) analysis [50] was used to identify differentially abundant microbial taxa between control and CD at all taxonomic levels. Relative abundance of taxon was calculated at each taxonomic rank level, namely kingdom, phylum, class, order, family, genus and species. For LEfSe analysis, all values were multiplied by 1000,000 according to the user instructions and the taxa with a logarithmic LDA score >2.0 were considered to be significantly different between groups.

### Multivariate and statistical analysis

PCA and PLS-DA were performed on quantified protein groups for unenriched samples and Kac sites for enriched samples. Briefly, the protein groups or Kac sites that were quantified in >50% of the samples were used and missing values were imputed using K-nearest neighbour (KNN) method (K=5). PCA and KNN imputation were performed in MATLAB (The MathWorks Inc.). PLS-DA was performed using MetaboAnalyst 4.0 [51] and the proteins or Kac sites that had a Variable Importance in Projection (VIP) value >2.0 were considered to be significantly different between control and CD groups.

Statistical significance of the difference between groups was evaluated using Mann-Whitney U test, unless otherwise indicated. The heatmap was generated using iMetaLab (http://imetalab.ca/). The Sankey plot was generated using SankeyMATIC (http://sankeymatic.com/).

## Availability of data and materials

All MS proteomics data that support the findings of this study have been deposited to the ProteomeXchange Consortium (http://www.proteomexchange.org) with the dataset identifier PXD013427 (http://proteomecentral.proteomexchange.org/cgi/GetDataset?ID=PXD013427).

## Acknowledgements

This work was supported by funding from the Natural Sciences and Engineering Research Council of Canada (NSERC), the Government of Canada through Genome Canada and the Ontario Genomics Institute (OGI-114 & OGI-149), CIHR grant number GPH-129340 and MOP-114872. DF acknowledges a Distinguished Research Chair from the University of Ottawa. We acknowledge Ruth Singleton and Christine Figeys at the CHEO, Ottawa ON, for their help in collecting intestinal aspirate samples. We also thank Dr. Kendra Hodgkinson at the University of Ottawa for her help in editing the manuscript.

## Authors’ contributions

D.F., A.S., D.M. and X.Z. designed the study. D.M. collected patient samples and clinical data. S.A.D. and K.W. pre-processed the samples. X.Z. and Z.N. performed the experiments and data analysis. M.S. and C.F. provided materials and involved in discussion of the study design. X.Z., D.F., A.S., D.M. and J.M. wrote the manuscript. All authors participated in the data interpretation, discussion and edits of the manuscript.

