## Supplementary Note and Figures for "Deep characterization of the protein lysine acetylation in human gut microbiome and its alterations in patients with Crohn’s disease"

Zhang *et al*.

### Supplementary Note 1

We performed taxonomic analysis using all quantified Kac microbial peptides and identified 3 kingdoms, 6 phyla, 10 classes, 10 orders, 19 families, 28 genera and 29 species with a minimum of 3 distinctive peptides (Supplementary Data 7). Linear discriminant analysis effect size (LEfSe) analysis showed that the acetylome-based abundances of species *Roseburia inulinivorans*, *Eubacterium eligens* and *Megamonas funiformis* were the most significantly decreased in CD compared to that of controls (Supplementary Fig. S3a). The acetylome-based abundance of *Bacilli* was the only one that was identified as significantly increased in CD compared to controls (Supplementary Fig. S3a). We also performed LEfSe analysis for metaproteome-based taxonomic compositions, which identified 12 taxa that were decreased and 10 taxa that were increased in CD compared to control subjects (Supplementary Fig. S3b). Interestingly, we found that the metaproteome of *Bacilli* was decreased in CD, which is opposite to the observations using the lysine acetylome-based abundances (Supplementary Fig. S3c). As shown in Supplementary Fig. S3d, the ratios of acetylome-based abundance to metaproteome-based abundance of *Bacilli* were significantly increased in CD compared to controls (P < 0.0001).

To further identify the taxa that exhibited significantly changed acetylome-to-metaproteome ratios, we performed LEfSe analysis using the ratios of all 103 taxa identified in lysine acetylome data set (phylum Chloroflexi and species *Roseburia intestinalis* were removed for this analysis due to detection levels below threshold for analysis in the metaproteome) (Supplementary Fig. S3e). The results showed that, in addition to *Bacilli*, *Ruminococcus* also exhibited significantly increased acetylome-to-metaproteome ratios in CD compared to control. The 6 taxa that showed increased acetylome-based abundances in CD compared to control subjects also exhibited significantly decreased acetylome-to-metaproteome ratios in CD.

### Supplementary Figures

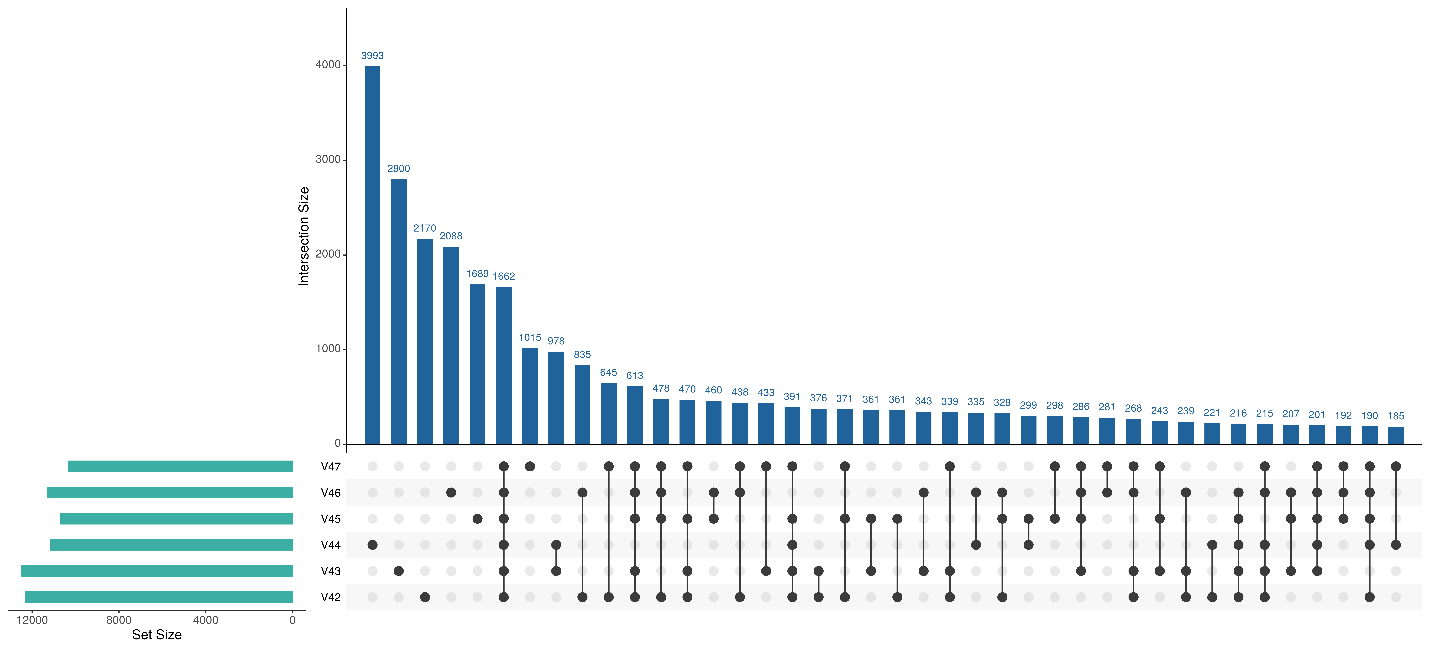

**Supplementary Fig. S1** Up set plot showing the overlap of quantified Kac sites among all six samples. Up set plot was generated using iMetaLab (<https://imetalab.ca/>).

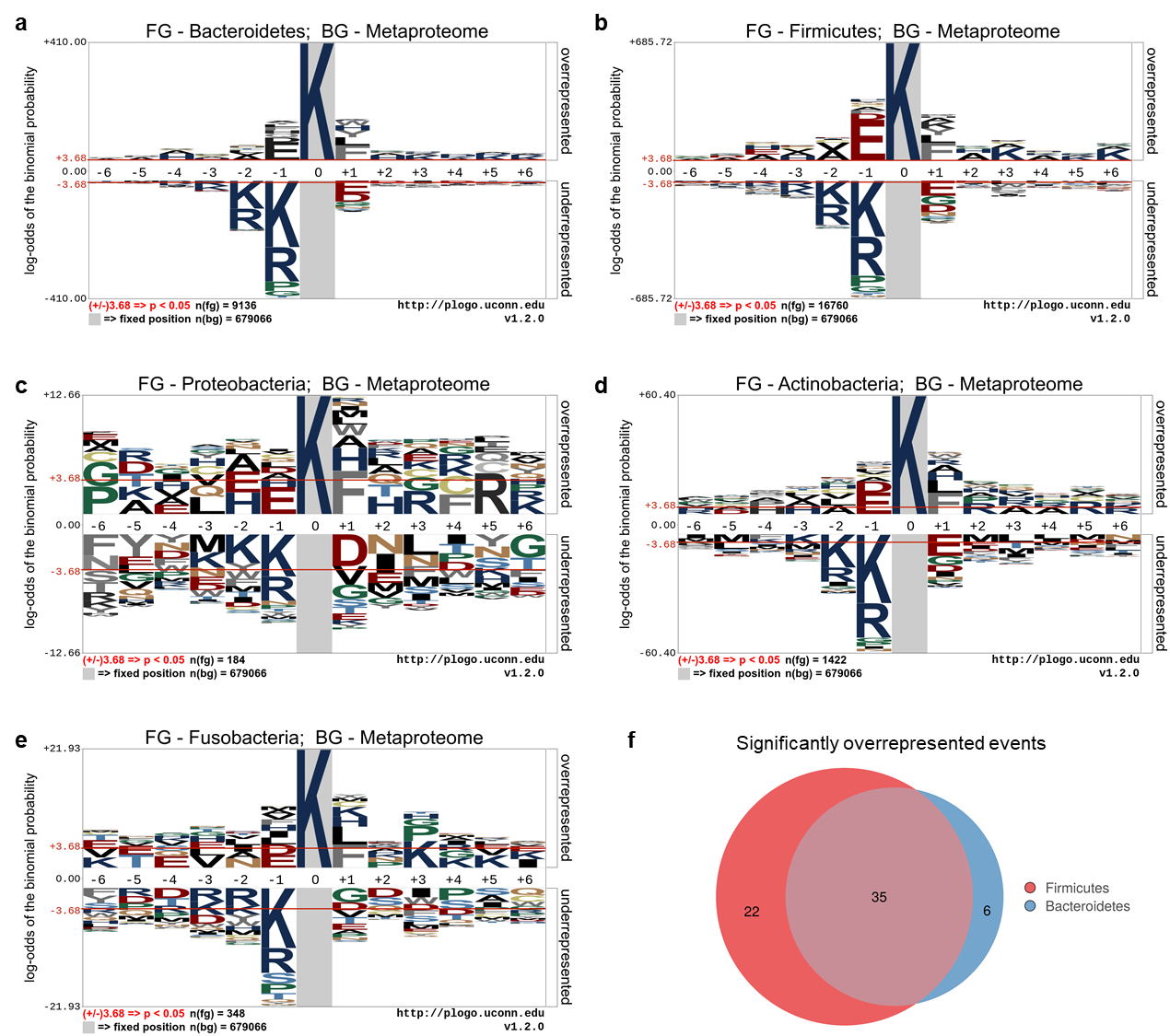

**Supplementary Fig. S2** pLogo analysis of phylum-specific Kac sites. (a-e) pLogo sequence visualizations of Bacteroidetes, Firmicutes, Proteobacteria, Actinobacteria, and Fusobacteria, respectively. (f) Overlap between the significantly over-represented positions of Firmicutes- and Bacteroidetes-specific Kac sites. The n(fg) and n(bg) values indicate the number of foreground and background sequences, respectively. The red horizontal bars on the pLogo correspond to a threshold of p < 0.05.

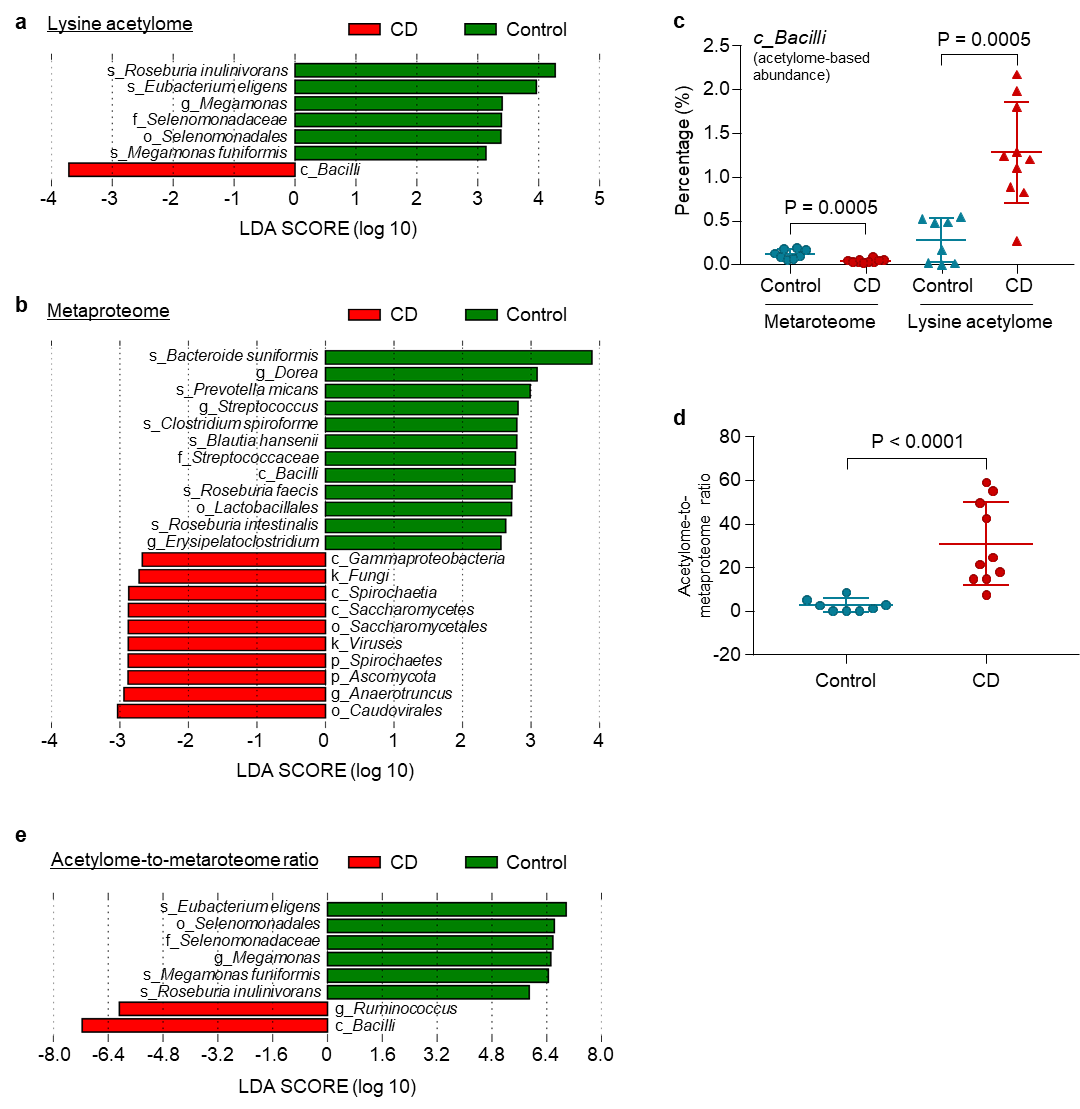

**Supplementary Fig. S3** Taxonomic alterations of protein acetylation in the pediatric CD microbiome. (a) LEfSe analysis of lysine acetylome-based taxonomic compositions; (b) LEfSe analysis of metaproteome-based taxonomic compositions; (c) Percentage of *Bacilli* in metaproteome and lysine acetylome data sets; (d) Acetylome-to-metaproteome ratios of *Bacilli* in pediatric CD and control subjects; (e) LEfSe analysis of the acetylome-to-metaproteome ratios of all quantified taxa in the lysine acetylome data set.

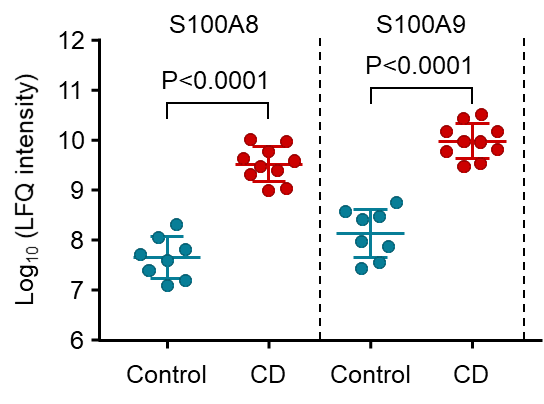

**Supplementary Fig. S4** Log_10_-transformed LFQ intensity of S100A8 and S100A9 in metaproteomic data set.

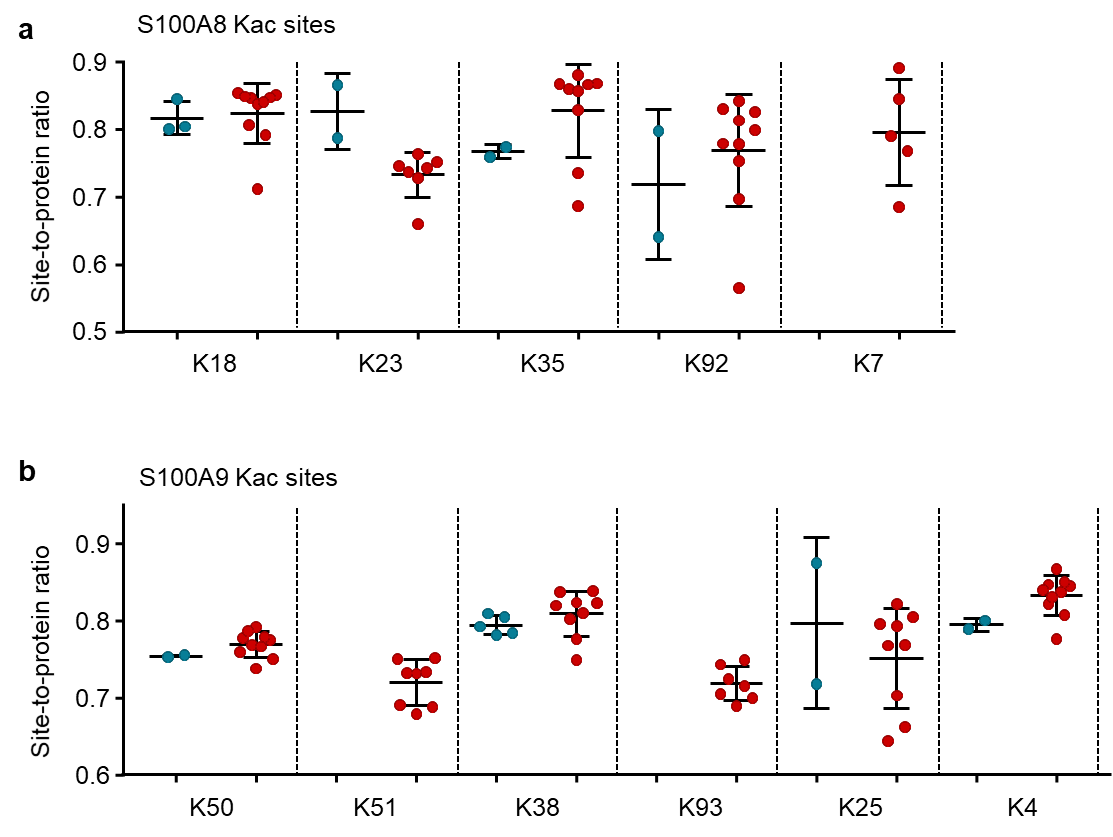

**Supplementary Fig. S5** Kac site-to-protein ratios of S100A8 (a) and S100A9 (b) in the intestinal aspirate samples of pediatric CD patients.
